# NetActivity enhances transcriptional signals by combining gene expression into robust gene set activity scores through interpretable autoencoders

**DOI:** 10.1101/2023.07.31.551238

**Authors:** Carlos Ruiz-Arenas, Irene Marín-Goñi, Liewei Wang, Idoia Ochoa, Luis A Pérez-Jurado, Mikel Hernaez

## Abstract

Grouping gene expression into gene set activity scores (GSAS) provides better biological insights than studying individual genes. However, existing gene set projection methods cannot return representative, robust, and interpretable GSAS. We developed *NetActivity*, a framework based on a sparsely-connected autoencoder and a three-tier training that yields robust and interpretable GSAS. *NetActivity* was trained with 1,518 well-known gene sets and all GTEx samples, returning GSAS representative of the original transcriptome and assigning higher importance to more biologically relevant genes. Moreover, *NetActivity* returns GSAS with a more consistent definition than GSVA and hipathia, state-of-the-art gene set projection methods. Finally, *NetActivity* enables combining bulk RNA-seq and microarray datasets in a meta-analysis of prostate cancer progression, highlighting gene sets related to cell division. When applied to metastatic prostate cancer, gene sets associated with cancer progression were also altered due to drug resistance, while a classical enrichment analysis identified gene sets irrelevant to the phenotype.

## Introduction

Although gene expression analyses have greatly improved our understanding of the physiopathology of multiple diseases and conditions, gene expression analyses performed at the gene level can be difficult to interpret, particularly when hundreds of genes are identified as differentially expressed, or when differentially expressed genes have an unknown function. In addition, measures of the same gene with different technologies (such as RNA-seq and microarrays) may present a reduced correlation^1^, resulting in different genes detected as differentially expressed^2^. Combining gene expression measurements into gene set activity scores (GSAS) have shown to address these critical issues^3,4^. To perform gene expression analyses, GSAS should have three important properties: (1) **representativeness** - GSAS should properly encode the transcriptional variance of the dataset; (2) **robustness** - biological insights provided by the GSAS should not change if GSAS are recomputed, and GSAS computed on similar samples using different technologies should be highly correlated; and (3) **interpretability** - researchers should be able to know which genes have higher importance in GSAS calculation.

Methods to project individual gene expression values into GSAS either prioritize their robustness and interpretability or their representativity. Methods that prioritize robustness and interpretability employ a weighted sum of the expression of the genes within the gene set. These weights can be uniform for all genes^5^, or assign positive and negative weights to genes based on the literature^6,7^. Fixing the gene weights ensures that GSAS definition remains consistent across datasets and the contributing genes are known. However, these methods lack representativity, as the gene weights are not proportional to the genes’ significance within the gene set. Alternative methods have been developed to model GSAS so they effectively capture the variability of the transcriptome. These methods include utilizing gene ranks^8,9^, maximizing the variability of genes within a gene set^10^, incorporating topological information from gene-gene co-expression networks^11,12^, or modeling the propagation of signals within pathways^13^. As the GSAS’ representativity is maximized within each dataset, it may lead to potential differences in the most relevant genes across datasets due to overfitting, compromising the GSAS robustness. Further, they present challenges in terms of interpretation, as researchers cannot readily determine which specific genes are more relevant for GSAS computation.

Shallow sparsely-connected autoencoders pose as ideal alternatives to address the shortcomings of current gene set projection methods^14–16^. Autoencoders are neural networks (NN) where the input data is reduced to a lower dimension and then expanded to reconstruct the original input data. Recently, shallow sparsely-connected autoencoders have been proposed to define GSAS^14–16^, where each gene set is represented by a neuron of the inner layer and is only connected to the genes in the gene set. The model aims to learn the low-dimensional embedding (i.e., the set of gene set scores) that best represents the input data. These methods also yield highly representative and interpretable GSAS, as GSAS are a weighted sum of the genes in the gene set. Nonetheless, current approaches generate GSAS heavily depending on the parameters’ initialization, and hence they are not robust.

In this work, we propose a computational framework, *NetActivity*, to define highly representative, robust, and interpretable gene set activity scores based on shallow sparsely-connected autoencoders. We trained the model by selecting gene sets from Gene Ontology (GO)^17,18^ Biological Processes and KEGG^19–21^ pathways and using the entire GTEx project^22^ data, showing that *NetActivity* generates GSAS independent of the initialization parameters that translate to unseen datasets representing different conditions. Further, *NetActivity* returns an importance score for each gene that agrees with their biological relevance in the studied context. The main model is distributed in a Bioconductor package to facilitate the computation of GSAS on new datasets. We compared *NetActivity* with *GSVA* and *hipathia*, two widely used state-of-the-art gene set projection methods, and found that *NetActivity* had the highest overall performance. Finally, we demonstrated some applications of *NetActivity* by (1) performing a meta-analysis in prostate cancer combining datasets from three different gene expression platforms; and (2) finding new biological insights in a metastatic castration-resistant prostate cancer dataset.

## Results

### *NetActivity* framework and main model

We have developed *NetActivity*, a framework to define sample-wise gene set activity scores (GSAS) from gene expression data. *NetActivity* consists of an autoencoder-like neural network (NN) that learns a low-dimensional representation of the gene expression data. The encoder maps the input gene expression vector *x ξ R^n^* onto a *p*-dimensional embedded representation (with *n ≥ p*). Each neuron on the embedded space represents a known biological gene set, so each neuron is only connected to the genes of this gene set (Figure 1A). GSAS are defined as the output of the embedded code before the activation function. Finally, the decoder is composed of a fully connected layer that aims to reconstruct the input gene expression vector from the GSAS (Figure 1A).

**Figure 1:**
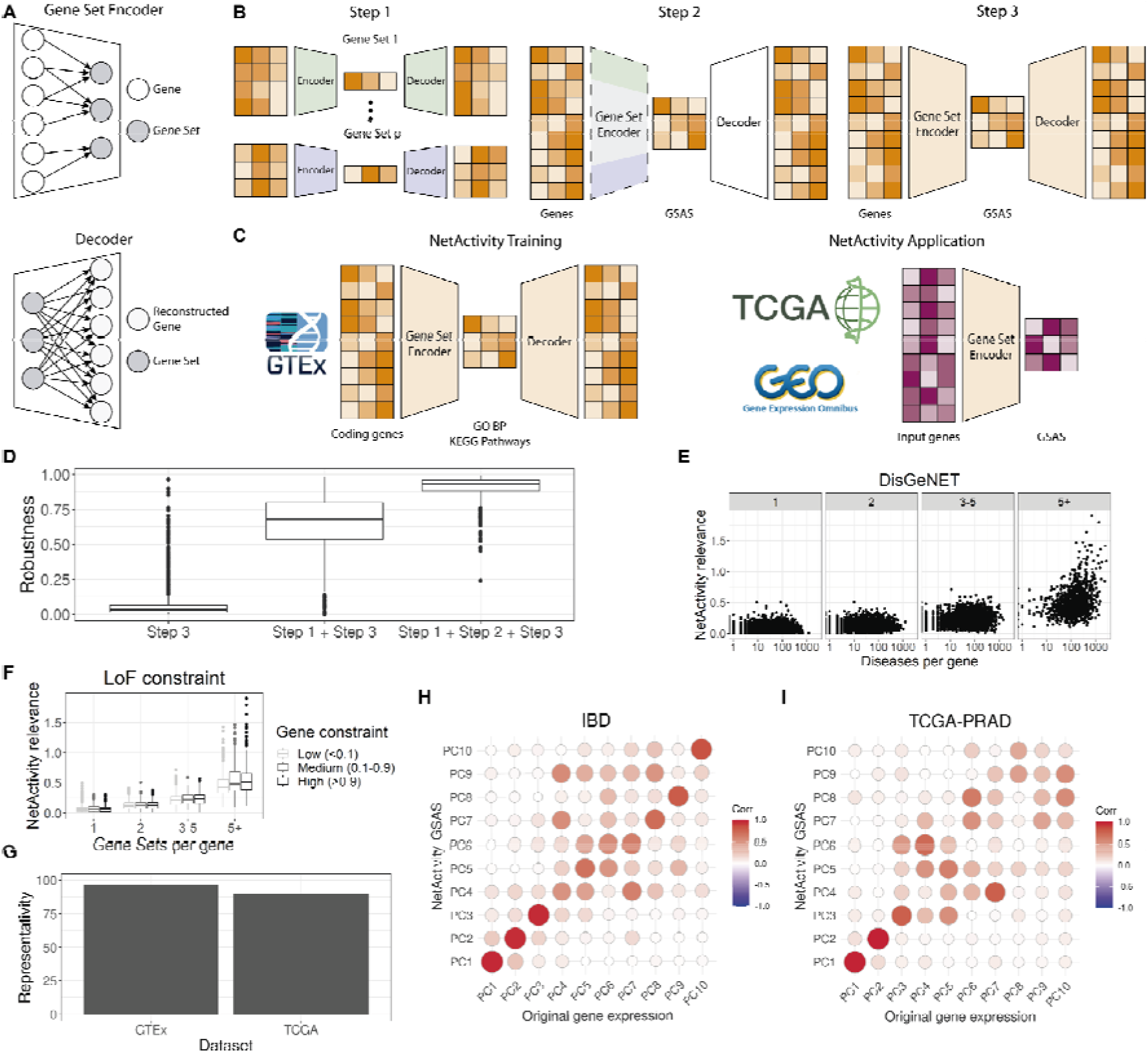
*NetActivity* framework to define gene set activity scores. **A:** Encoding and decoding layers. The gene set encoder layer encodes information from genes to gene sets. Each gene set neuron only receives information from the genes present in the gene set. The decoder layer aims to reconstruct the original gene expression matrix by combining the information from all gene sets via a fully connected layer. *NetActivity* framework enables to include multiple layers in the gene set encoder and decoder. **B**: *NetActivity* three-step training. In step 1, an autoencoder is trained for each gene set. In step 2, weights from the encoders of step 1 are used to initialize the gene set encoder layer. The weights of this layer are frozen (represented by dashed lines), while the decoder is trained. In step 3, the whole network is trained. GSAS: Gene Set Activity Scores. **C**: Definition of the main model. The main model is trained using GTEx data and includes GO (Gene Ontology) Biological Processes (BP) and KEGG Pathways as gene sets. The trained gene set encoder is used to compute GSAS in other datasets. **D**: Robustness of model training. Robustness measures how robust each GSAS is to parameters initialization. A 0 means that the GSAS completely depends on the parameter initialization while a 1 means that is completely independent. We evaluated the robustness of three training processes: Step 3, i.e., directly training the whole autoencoder; Step 1 + Step 3, i.e., training the whole autoencoder after initializing the parameters by running an autoencoder in each gene set; Step 1 + Step 2 + Step 3, i.e., using the three-steps training. **E-F**: Biological relevance of *NetActivity* weights. *NetActivity* gene relevance estimates the importance of a gene for defining GSAS. The higher the gene relevance, the more gene sets a gene has a strong influence on. **E**: Correlation between *NetActivity* gene relevance and number of diseases associated with a gene. Genes are grouped by the number of gene sets they belong to (1, 2, 3-5, 5+). X-axis represents the number of diseases a given gene is associated with in the DisGeNET database^24^. **F**: Correlation between *NetActivity* gene relevance and loss of function intolerance (pLI). Gene Sets per gene: number of gene sets where a gene is included. Gene constraint: a measure of the intolerance of a gene to loss of function mutations. Gene constraint categories are computed based on pLI scores from gnomAD: low (<0.1), medium (0.1-0.9), and high (>0.9). A higher pLI means a stronger depletion of loss of function mutations and higher biological relevance. **G**: Representativity of the GSAS. Representativity measures the proportion of variance of the original gene expression matrix contained in the whole GSAS matrix. Representativity was measured in the dataset used to train the data (GTEx) and in an external dataset (TCGA). **H-I**: correlation between the top 10 principal components (PCs) of the gene expression matrix and the GSAS matrix in an IBD cohort (H) and the tumor cohort PRAD from TCGA (I).

*NetActivity’s* architecture ensures that GSAS are interpretable, while our three-step training ensures that GSAS are representative and robust. In step 1, we initialize the encoder layer of *NetActivity* by training an autoencoder for each gene set, with the genes in the gene set as input features and one neuron in the embedding layer (Figure 1B). This step learns the weights so that the output of the embedding layer maximizes the representativity of the expression of the genes in the gene set. In step 2, we freeze the weights from the encoder layer and train the decoder. This step ensures that GSAS are independent of the initialization parameters and hence, robust, by ensuring a smooth training, as done in transfer learning^23^. In step 3, we unfreeze all the weights and fine-tune the network by training for additional epochs (Figure 1B). At the end of the training, the set of all GSAS will accurately encode the input gene expression, hence optimizing the representativity of the whole transcriptional variability.

*NetActivity’s* main model had one layer on both the encoder and decoder. Our main model contained a selection of 1,518 well-known biological gene sets (Sup Table 1): 1,485 (Gene Ontology) Biological Processes^17,18^ (BP) (Sup Table 2) and 33 KEGG^19–21^ Pathways (Sup Table 3). The model was trained using all GTEx project^22^ samples (9,662), so the model defined GSAS representative of multiple tissues and biological functions (Figure 1C). This selection of gene sets in combination with the three-step training returned highly robust GSAS, i.e. independent of the weights’ initialization (Figure 1D). In contrast, including all GO terms and KEGG pathways reduced the GSAS robustness (Sup Fig 1). While *NetActivity* enables adding additional hidden layers or dropout, none of these configurations significantly improved the performance to compensate for the increased complexity and reduced interpretability of the model (Sup Figs 2-5). Finally, this model is openly distributed in a Bioconductor package (www.bioconductor.org/packages/NetActivity) to facilitate the computation of GSAS on new datasets (Figure 1C).

**Figure 2:**
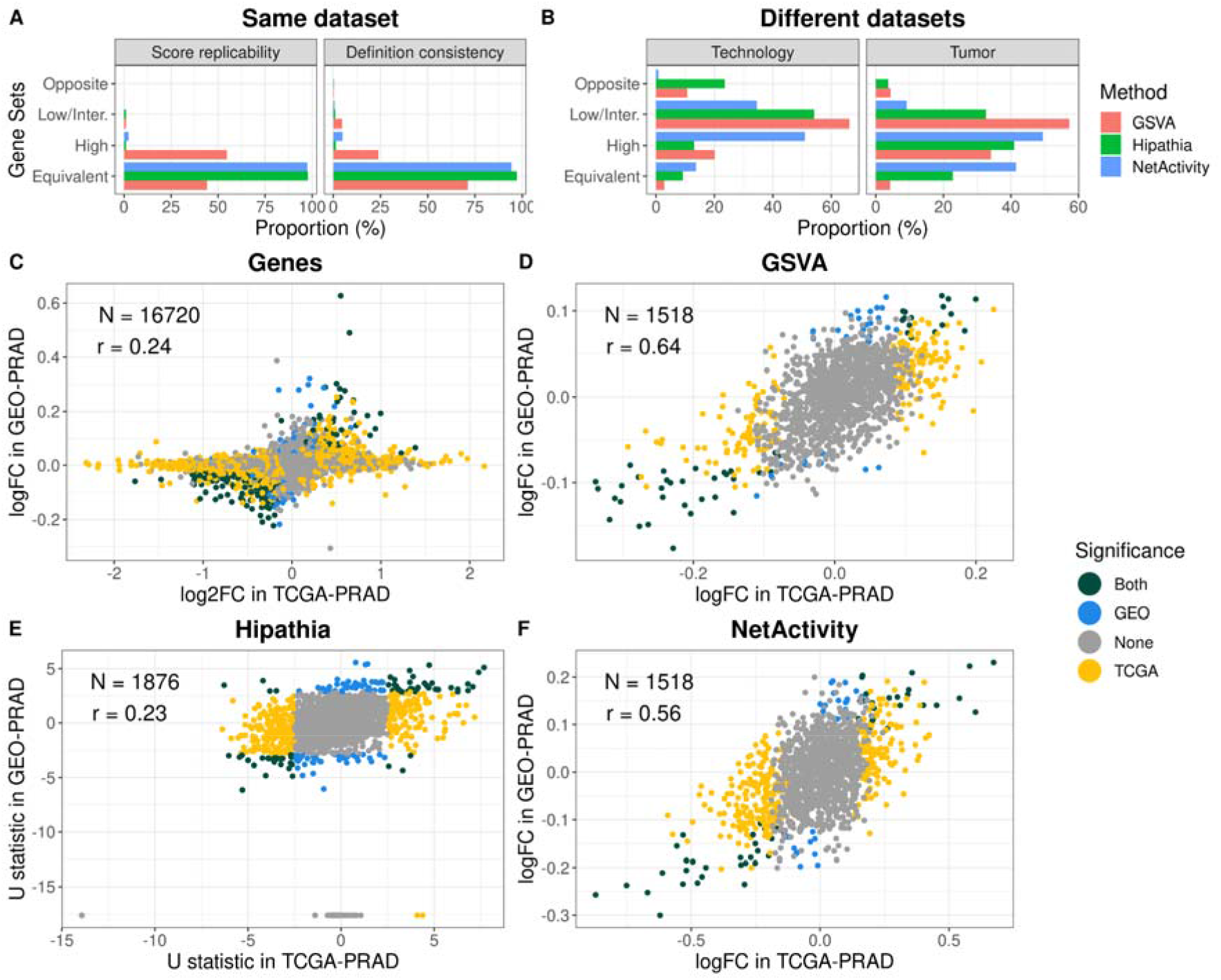
Comparison of gene set projection methods. **A-B**: Consistency of GSAS definitions. Equivalent: value > 0.9. High: value > 0.7. Low/Inter.: value > 0. Opposite: value < 0. **A**: Consistency of gene set score definitions depending on the overall population. For the three methods, we computed gene set scores on control samples from TCGA-PRAD. We computed the scores using two populations: (1) all PRAD samples; and (2) control PRAD samples. Score replicability: Pearson correlation between the gene set scores computed in populations 1 and 2. Definition consistency: we defined the definition of a gene set score as the Pearson correlation between each gene and the gene set score it belongs to. Definition consistency was the Pearson correlation between the gene set definitions in populations 1 and 2. **B**: Consistency of gene set score definitions in different datasets. Technology: consistency of the gene set score definition in two datasets from prostate cancer generated with RNA-seq and gene expression microarray. Tissue: consistency of the gene set score definition in two datasets from prostate cancer and breast cancer generated with RNA-seq. **C-F**: Differences in gene expression between Gleason low and high using 4 approaches and 2 datasets (TCGA-PRAD: PRAD samples; GEO-PRAD: GSE169038). **C**: Differential expression of genes. logFC: differential expression estimates from *limma*. log2FC: differential expression estimates from *DESeq2*. **D**: Differential expression of GSVA scores. **E**: Differential expression of hipathia pathways. Axes report the U statistic of the Wilcoxon test. **F**: Differential expression of *NetActivity* GSAS. LogFC is the differential expression estimate from *limma*. Gene sets (or genes) were colored by their statistical significance in the datasets: Both (green) - features with different expression in both datasets (FDR < 0.05); TCGA (yellow) - features with different expression in TCGA-PRAD (FDR < 0.05) but not in GEO-PRAD (FDR > 0.05); GEO (blue) - features with different expression in GEO-PRAD (FDR < 0.05) but not TCGA-PRAD (FDR > 0.05); None (grey) - features without differences of expression in any of the datasets (FDR > 0.05).

### NetActivity’s three-tier training produces interpretable and biologically consistent GSAS

We evaluated whether *NetActivity* produced biologically relevant and interpretable GSAS. *NetActivity* gene relevance was defined as the sum of the genes’ weights magnitudes across multiple gene sets (see Methods). We compared the inferred *NetActivity* gene relevance with two measures of biological relevance: (1) the number of diseases associated with a gene in DisGeNET, a database of gene-disease associations^25^; and (2) the intolerance to loss of function mutations (pLI) from gnomAD^26^. Genes with a high intolerance to loss of function mutations are likely to be essential for normal biological function. Genes with higher *NetActivity* relevance were associated with more diseases (Figure 1E, p-value < 2e-16) and were more intolerant to loss of function mutation (Figure 1F, p-value = 1.1e-5), after correcting for the number of gene sets a gene is included into. These results support that *NetActivity* gene relevance is correlated with biological gene relevance.

To delve into the learning process of *NetActivity*, we investigated the gene set weights and the corresponding GSAS for the hsa00430 KEGG pathway (taurine and hypotaurine metabolism) (Sup Figure 6) at the three training steps. In step 1, the two main clusters of correlated genes were initialized with the largest weights (Sup Fig 7-8), indicating that weights are capturing the largest sources of variance (i.e., information) in the data. Further, the model converged to similar weights across different parameter initializations (Sup Fig 7), due to the much lower number of parameters (between 10 and 30, corresponding to the number of genes in a gene set) with respect to the 9,662 samples of GTEx. Step 2 ensures that gene weights are robust after step 3 (Sup Fig 7), as it avoids losing the information stored in the encoder while the decoder weights are transitioning from their random initialization^23^. Finally, step 3 refined the encoder and decoder weights to obtain a biologically meaningful representation of the gene set. On one hand, step 3 reduced the weights of GAD1 and GAD2 to close to 0 (Sup Fig 9). While the role of GAD1 and GAD2 in the taurine/hypotaurine metabolism in humans is not well supported, both genes were expressed almost exclusively in the brain (Sup Fig 10), representing a relevant source of the gene set variance. On the other hand, step 3 further increased the weight of FMO1 (Flavin containing monooxygenase 1)(Sup Fig 9), the main gene involved in the catalysis of hypotaurine to taurine reaction^27^. Therefore, step 3 increased the weights of genes relevant to the gene set function, while reducing the weights of irrelevant genes, although they contribute to the gene set variance.

GSAS from *NetActivity* were also representative of the whole transcriptome, retaining most of the transcriptional variance of the input samples, both in the training dataset (GTEx) and in a new dataset of cancer samples (TCGA) (Figure 1G). Further, principal components (PCs) of GSAS explained a similar proportion of variance to PCs of the input gene matrix (r = 0.99), both in an external common disease cohort (IBD cohort) or in each TCGA tumor subtype (cancer). Indeed, the first 10 PCs of the original gene matrix were highly correlated with one of the first 10 PCs of the GSAS matrix in both cases (Figure 1H-I).

### *NetActivity*, unlike previous methods, yields robust GSAS across different technologies and datasets

We compared the robustness of *NetActivity* GSAS with two state-of-the-art methods to encode GSAS: GSVA^9^ and *Hipathia*^13^. First, we evaluated the consistency of the GSAS definition across datasets, defined as the correlation between the GSAS and the expression of the genes in the gene set (see Methods). Thus, a method would return consistent GSAS definitions if the GSAS computed in different datasets are correlated with the same genes. We explored the consistency of the GSAS definition in: (1) subsamples in a dataset; (2) datasets generated with different technologies; and (3) datasets from different tissues.

We computed GSAS using all TCGA-PRAD samples (prostate cancer project from TCGA) or only the subset of healthy tissue samples, and compared the GSAS of healthy samples in both approaches. *NetActivity* and *Hipathia* returned equivalent GSAS (i.e. GSAS whose Pearson correlation > 0.9) and consistent GSAS definitions for more than 90% of the gene sets (Figure 2A). In contrast, only 71% of GSVA GSAS had equivalent values and 44% consistent definitions. Next, we compared GSAS definitions in TCGA-PRAD with those obtained in a dataset from: (1) the same tissue (prostate cancer) but generated with another technique (gene expression microarray samples from GSE169038, GEO-PRAD); or (2) a different tissue (breast cancer, TCGA-BRCA) generated with the same technique (RNA-seq). In both cases, *NetActivity* returned the most consistent GSAS definitions (Figure 2B), having the highest proportion of gene sets with a highly consistent definition. In GSVA most of the gene sets had different GSAS definitions, either when comparing different technologies or different tissues.

Second, we evaluated the biological replicability of the GSAS. We first performed a differential expression analysis between samples with low or high Gleason score, a histologic measure of tumor progression, in: (1) PRAD samples from TCGA, generated with RNA-seq (TCGA-PRAD); and (2) prostate cancer samples from GSE169038, generated with a gene expression array (GEO-PRAD). We evaluated the concordance of the results when using single gene expression analysis or the gene set projection methods (*NetActivity*, *GSVA,* and *Hipathia*) (Figure 2C-F). *Hipathia* presented the lowest reproducibility (Figure 2E), even with gene sets significant in both datasets but in opposite directions; while *NetActivity* and *GSVA* had the most replicable results (Figure 2D and 2F). For both methods, all gene sets differentially expressed in both datasets had a concordant direction. However, gene sets differentially expressed in *GSVA* had a modest overlap with the gene sets differentially expressed with *NetActivity* (Sup Fig 11) because *GSVA* was limited to gene sets where genes had coherent differences between the groups (Sup Fig 12). Notice that differential expression using individual gene expression measurements was noisier, with some genes presenting large gene expression differences in one dataset but not in the other (Figure 2C), probably due to technological differences.

### *NetActivity* enables performing an interpretable gene set meta-analysis combining multiple gene expression platforms

We run a GSAS meta-analysis on differences between Gleason low and Gleason high prostate cancer samples. We included nine prostate cancer datasets: three generated with RNA-Seq, three with Illumina HumanHT-12 array, and three with the Affymetrix HuEx1.0 array (Sup Table 4). 152 gene sets had differential GSAS between Gleason low and high samples (FDR < 0.05). Several of them were associated with cell division and morphogenesis (Sup Table 5), coherent with differences in tumor progression between Gleason low and high. The magnitude of the GSAS was comparable between the different technologies (Figure 3), which is essential to enable a meta-analysis.

**Figure 3:**
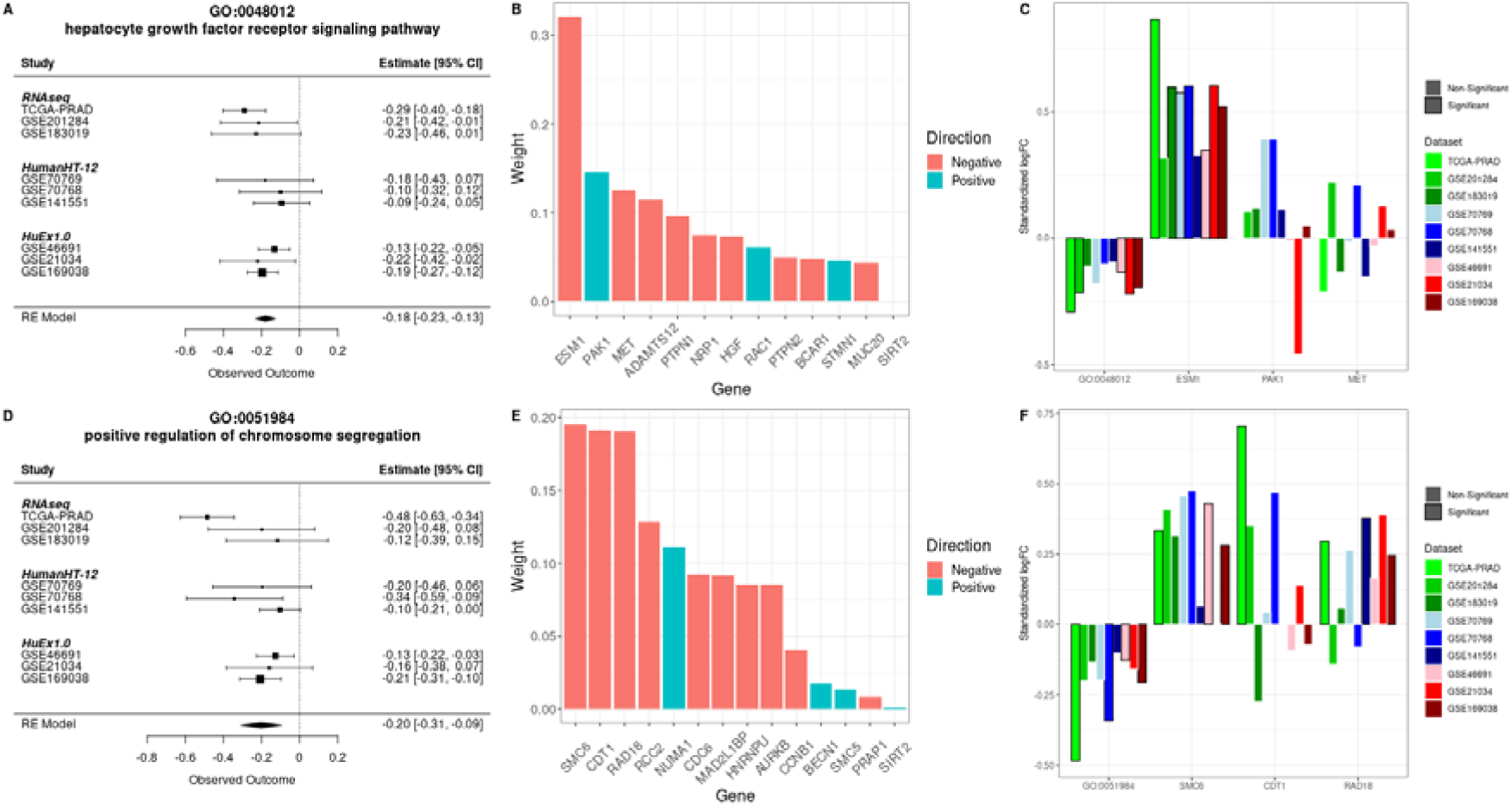
Gene set meta-analysis in prostate cancer. Differences in gene set scores between Gleason low and high samples were analyzed in nine cohorts. **A-C**: Analysis of GO:0048012 gene set. A: Meta-analysis of GO:0048012 gene set. Forest plot shows the effects’ magnitude and their standard error in the 9 datasets. **B**: Weights for GO:0048012 in *NetActivity*. Weights are in absolute value to compare their magnitude. **C**: Difference in expression in high Gleason samples with respect to low Gleason samples. Gene expression values were standardized before running the analyses, so the coefficients are comparable. Bars highlighted in black represent associations nominally significant (p-value < 0.05). **D-F**: Analysis of GO:0051984 gene set. **D**: Meta-analysis of GO: 0051984 gene set. **E**: Weights for GO:0051984 in *NetActivity*. **F**: Difference in expression in high Gleason samples with respect to low Gleason samples. Gene expression values were standardized before running the analyses, so the coefficients are comparable. Bars highlighted in black represent associations nominally significant (p-value < 0.05). LogFC is the differential expression estimate from limma.

Further, we explored the two top gene sets associated with Gleason scores: GO:0048012 (hepatocyte growth factor receptor signaling pathway) and GO:0051984 (regulation of chromosome segregation). Individuals with high Gleason scores had lower GSAS of the hepatocyte growth factor receptor signaling pathway, a well-known pathway of prostate cancer progression^28^ (Figure 3A). This GSAS is mainly driven by *ESM1*, which has negative weights indicating the negative correlation of *ESM1* expression with the activity of the hepatocyte growth factor receptor signaling pathway (Figure 3B). Thus, individuals with high Gleason scores had higher expression of *ESM1* (Figure 3C). Similarly, individuals with high Gleason scores had lower GSAS of positive regulation of chromosome segregation (Figure 3D). In this case, three genes (*SMC6*, *CDT1,* and *RAD18*) drive this gene set’s GSAS, the three with negative weights (Figure 3E). In general, individuals with high Gleason had higher expression of these genes (Figure 3F), although individual gene expression had more variable differences than the GSAS.

### NetActivity advances the knowledge of prostate cancer

We applied *NetActivity* to the PROMOTE study^29,30^ to show its performance in a clinical dataset. The PROMOTE study consists of bone metastatic samples from metastatic castration-resistant prostate cancer patients treated with abiraterone. We characterized whether differences in gene expression were associated with a better response to abiraterone (higher Time To Treatment Change or TTTC), using *NetActivity* and a traditional differential expression analysis. 30 gene sets had GSAS associated with response to treatment (FDR < 0.05), with top gene sets related to cell division and cell cycle processes (Sup Table 6). Interestingly, 19 top gene sets associated with response to treatment were also associated with tumor progression in the prostate cancer meta-analysis, suggesting that abiraterone treatment reverts cancer progression. Differential gene expression analyses revealed 592 genes differentially expressed (FDR < 0.05 and log2 fold-change > 1, Sup Table 7) and 248 enriched biological process GO terms (Sup Table 8, FDR < 0.05). In contrast, most of the top enriched terms were related to muscle tissue, hence irrelevant for the phenotype, while only a few were related to cell cycle and mitosis (Sup Fig 13). Even when restricting the analysis to genes and GO terms present in *NetActivity* (Sup Table 9), a high proportion of enriched terms were still related to muscle tissue.

*NetActivity* GSAS correlated better with response to treatment than individual genes. For instance, GSAS of the top significant gene set, regulation of attachment of spindle microtubules to kinetochore (GO:0051988), had a stronger association with TTTC (R2 = 0.41) than any of these genes individually (Sup Figure 14). In this gene set, samples with a better response (higher TTTC) had lower GSAS (Figure 4A). *NetActivity* assigned positive weights to the top three genes of this gene set (*KNSTRN*, *ECT2,* and *RCC2*) (Figure 4B), so these genes also had a negative correlation with TTTC (Figure 4C). Nonetheless, KNSTRN, ECT2, and RCC2 were not significantly associated with response to treatment after accounting for multiple testing, so these genes were not identified in the traditional gene expression analysis. Therefore, *NetActivity* can uncover relevant differences in key gene sets which might be missed in a traditional gene expression analysis.

**Figure 4:**
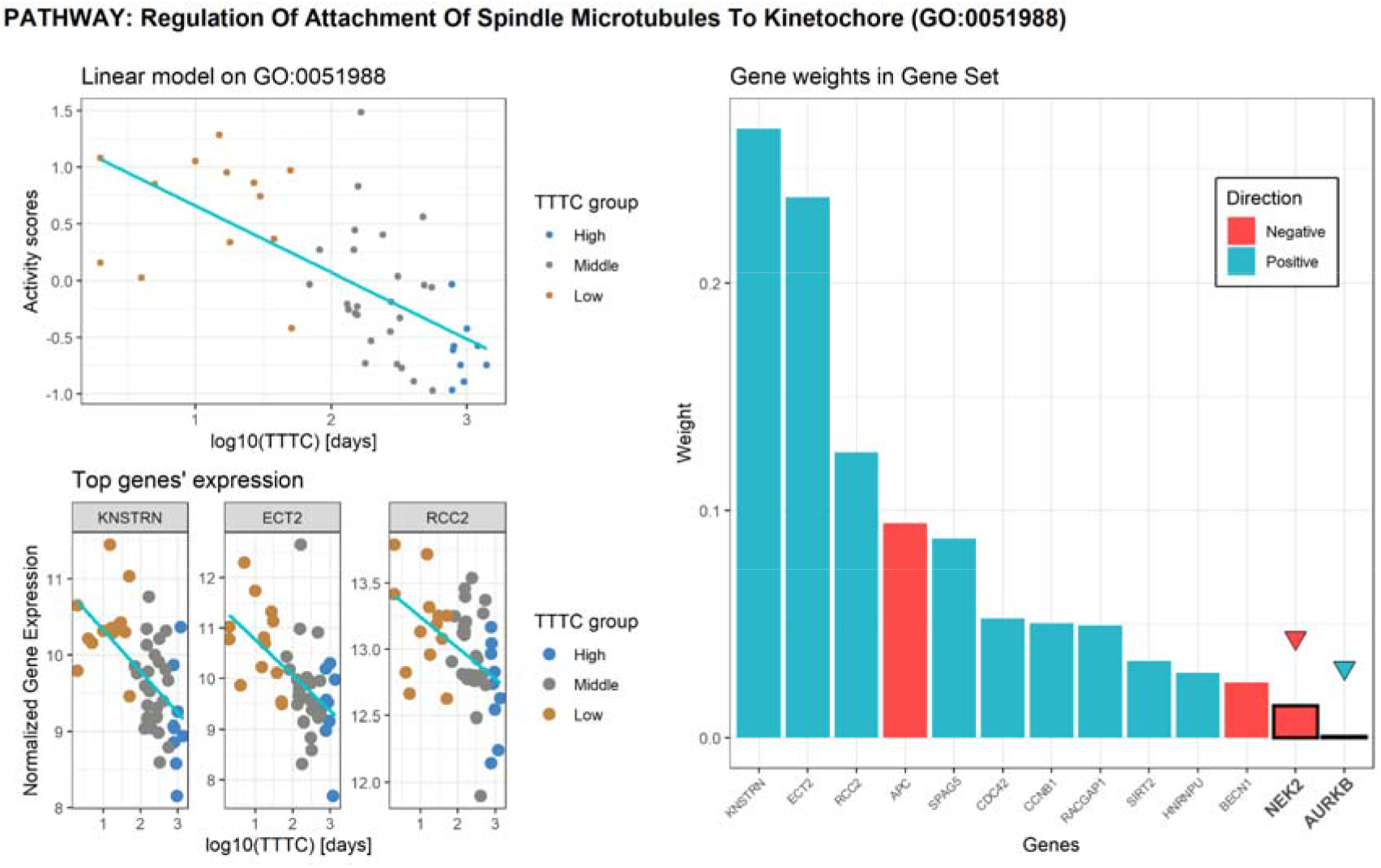
*NetActivity* application on PROMOTE study. **A**: NetActivity scores on GO:0051988 pathway by response phenotype (TTTC) and the fit of a simple linear regression model. **B**: *NetActivity* GO:0051988 weights; highlighted genes are differentially expressed on the *DESeq2* pipeline. C: normalized gene expression of top 3 genes of GO:0051988 by response phenotype with simple linear model regression on the TTTC. Patients are colored based on adjusted TTTC quartiles: ‘high’ - highest quartile so better response to therapy; ‘low’ - lowest quartile so considered as resistant to therapy; ‘medium’ – second and third quantiles.

### *NetActivity* framework is publicly available in Bioconductor and as a Nextflow pipeline

We have implemented the framework in *NetActivityTrain* (https://github.com/yocra3/NetActivityTrain), a Nextflow pipeline^31^ containing the steps for preprocessing the data, training the model, and applying it to new datasets. The model training is performed in Tensorflow - Keras 2.7^32^. *NetActivityTrain* outputs the model and the weights required for computing the gene set activity scores in new datasets and can be also used to compute gene set activity scores in new datasets. All the software and dependencies required to run *NetActivityTrain* are encapsulated in two docker images, publicly available in DockerHub (https://hub.docker.com/).

We have also developed a R/Bioconductor package, *NetActivity* (https://bioconductor.org/packages/NetActivity), that facilitates the computation of gene set activity scores in new datasets. The current implementation comes with the model trained in GTEx using selected GO biological processes terms and KEGG pathways. *NetActivity* can also standardize the data and manage genes present in the model but absent in the input data. *NetActivity* can also be used to compute the gene set activity scores of other models trained with the *NetActivityTrain*.

## Discussion

In this work, we propose *NetActivity*, a computational framework to define gene set activity scores (GSAS) from individual gene expression values using a sparse autoencoder-like neural network. *NetActivity* GSAS addresses three long-sought features in GSAS: they are representative of the input gene expression, robust to different initializations and technological biases, and interpretable such that biological insights can be inferred from the weights assigned to genes. Specifically, *NetActivity* first learns the best representation of each gene set in a reference dataset, assigning a weight for each gene in a gene set. Thus, *NetActivity* can accommodate gene sets with genes operating in opposite directions and give less weight to less relevant genes. In contrast, some gene set project methods, such as z-score^5^ or *singscore*^7^, assign the same weight to each gene in a gene set and are designed to detect gene sets with genes expressed in the same direction. Second, *NetActivity* defines robust GSAS, as the same weights are used to infer GSAS in new datasets. In contrast, gene set projection methods, such as z-score, *ssGSEA*^8^, *GSVA*^9^, *iPAS*^33^, *PLAGE*^10^, or *Pathifier*^34^, define gene weights based on the data, so the GSAS definition depends on the dataset where they are created. Thus, a high GSAS can be associated with high expression of a different set of genes depending on the dataset. Indeed, as shown for *GSVA*, the GSAS of a sample might depend on the samples included in the score computation. Third, although *NetActivity* starts by defining GSAS that maximize the variability of the genes in each gene set, the model can learn a gene set representation more representative of the gene set biology, especially when the gene set might not be accurately defined, as in the KEGG pathway hsa00430. Thus, gene set projection methods that maximize the variability of the genes in the gene set, such as *PLAGE* or *Pathifier,* might return gene set activity scores that do not represent the biological activity of the gene set. Fourth, the proposed framework allows training a model including any gene set, without additional biological knowledge. By contrast, methods such as *Oncofinder*^6^, *iPANDA*^11^, or *singscore*, require knowing which genes are positively or negatively associated with the gene set activity; while others, such as *TAPPA*^12^ or *Hipathia*^13^, require knowing the relationships between the genes in the gene set, restricting their application to well-defined pathways. Finally, once the model is trained and the gene set weights defined, *NetActivity* GSAS are computed individually for each sample. In contrast, some gene set projection methods, such as *Pathifier*, *iPAS,* or *GRAPE*^35^, require a reference panel to define the scores so they are not suitable for settings such as in epidemiology - where all samples are healthy and small differences are expected - or in some clinical trials - where all individuals are cases and present unique features.

*NetActivity* R/Bioconductor package implements our model trained with GTEx project^22^ data based on 1,518 GO^17,18^ biological process terms and KEGG^19–21^ pathways. This package enables users to compute GSAS in new datasets and currently is the best approach to define representative, robust, and interpretable GSAS from individual gene expression values. *NetActivity* generates scores with high replicability and independent of the population, can be applied to RNA-seq or gene expression array data, and has good biological reproducibility, as it can replicate the same biological results in datasets generated with different techniques.

*NetActivity* training can be used to gain insights into gene regulatory programs. Genes with low weights after training are considered less relevant for the gene set function. Thus, this information could be extracted from the model to remove less relevant genes and refine the gene sets. *NetActivity* also enables integrating different gene set collections in the same model. We trained our main model using GO biological processes terms and KEGG pathways, expecting to cover most of the human tissue functionalities. Nonetheless, *NetActivity* models can accommodate additional gene sets representing other biological functions^36^, gene regulatory networks^37^, disease transcriptomic signatures^36,38^, drug perturbation signatures^39,38^, or even custom gene sets. As shown in the main model, GSAS replicability can help us define a list of gene sets with independent biological functions. For example, if the model includes two gene sets sharing a high proportion of genes, they are very likely to represent the same unique biological function. As such, the corresponding GSAS will highly depend on the initialization of the weights. Based on this idea, we can remove gene sets until all included gene sets are robust, to identify an independent list of gene set functions (potentially from different collections). Finally, large gene sets had lower GSAS robustness, possibly because they do not represent a single biological function.

*NetActivity* advances in the integration of gene expression results and datasets obtained from different platforms. *NetActivity* returns gene set activity scores for 1,518 gene sets, independent of the platform; and the data preprocessing ensures that the GSAS are on the same scale. Thus, *NetActivity* facilitates performing large meta-analyses including datasets from different platforms, as we exemplified in a meta-analysis of nine public prostate cancer datasets generated with three platforms. Our meta-analysis demonstrated that GSAS reflect consistent biological processes that are associated with the phenotypes of interest. In our case, the top identified gene sets were relevant for prostate cancer progression, such as the hepatocyte growth factor receptor signaling pathway^28^. Hence *NetActivity* overcomes the current limitations of meta-analyses combining different gene expression platforms, such as having different features measured in different platforms^1^ or having the gene expression measured on different scales. *NetActivity* standardization achieves similar statistical properties to preprocessing gene expression data with quantile normalization and log2 transformation, an approach successfully applied to run a meta-analysis using effect estimates in Down Syndrome^40^. *NetActivity* also has some advantages over QuSAGE^41^, a method to perform meta-analysis based on gene set activity scores. As QuSAGE does not include any previous information about the relationships between genes, GSAS might represent overexpression of different genes in different datasets. In addition, QuSAGE has not been shown to be capable of integrating gene expression results from different platforms. Thus, *NetActivity* appears as the best-performing method to perform meta-analyses of multiple gene expression platforms.

*NetActivity* GSAS are also highly interpretable. Specifically, based on *NetActivity* weights, researchers can identify which genes are driving the GSAS and check the expression of these genes. Thus, in the meta-analyses of prostate cancer, we explored two top gene sets whose GSAS were driven by *ESM1*, and by *SMC6*, *CDT1,* and *RAD18*. Interestingly, *ESM1* and *CDT1* have been previously associated with prostate tumor progression^42,43^. Similarly, the top gene set whose GSAS was different between resistant and sensitive metastatic prostate samples to abiraterone treatment was driven by *KNSTRN*, *ECT2,* and *RCC2* genes, three genes previously associated with prostate cancer progression^44–46^. Therefore, *NetActivity* identifies gene sets associated with a condition, as well as the specific genes that drive this gene set scores, enabling researchers to conduct experimental validation of their results. This validation cannot be performed with commonly used gene set projection methods, such as GSVA, as they do not report the genes that drive the score computation.

In addition, *NetActivity* can improve the interpretability of standard gene expression analysis. In our application to metastatic-prostate cancer samples, *NetActivity* GSAS identified that abiraterone treatment modified gene sets associated with cell division and cell cycle. In addition, different gene sets presented similar differences in GSAS due to tumor progression (difference between Gleason high and low samples) or to abiraterone treatment, suggesting that effective abiraterone treatment reverses tumor progression. In contrast, traditional gene expression analysis identified gene sets related to muscle function, which are not relevant neither to the phenotype (metastatic prostate cancer) or to the tissue (bone). Two reasons can explain the better performance of *NetActivity* over traditional gene expression enrichment analysis. First, *NetActivity* can combine individual genes with small associations with the phenotype into a score, even if none of the individual genes pass the multiple testing corrections; whereas gene set enrichment analysis only considers genes passing the multiple testing correction. Second, *NetActivity* gives a different weight to each gene in the gene set, so relevant genes can have higher importance; while the traditional enrichment analysis gives the same importance to all genes. These features highlight the potential of *NetActivity* to get new biological insights from gene expression data.

*NetActivity* also has some limitations. First, GSAS sign is arbitrary and depends on the weights’ initialization. Thus, positive and negative values were considered equivalent, and all robustness measures were reported in absolute value. Nonetheless, exploring the magnitude and direction of the weights of the genes in the gene set enables us to understand what the GSAS represent. Second, the biological function of the gene set can be difficult to define. For instance, in gene metabolic pathways, a gene set can represent the production of different metabolites, which are not defined a priori. Therefore, further analyses and experiments are required to better characterize each gene set function. Finally, extending the model to new gene sets requires training the whole model in a large dataset. This requirement might not be feasible if we aim to define gene sets specific to rare conditions. One solution could be to fine-tune the model with the new dataset using techniques from transfer learning. More gene sets could also be included by adding new nodes in the encoder and initializing the weights with the pre-trained ones. Further studies are required to ascertain the validity of these approaches.

## Methods

### Gene set activation framework

*NetActivity* is based on an autoencoder-like neural network (NN). On its main configuration, *NetActivity* consists of an input layer (gene expression vector), a single encoder layer (gene set activity scores or GSAS), and a decoder layer (reconstructed gene expression vector). The network is trained in three steps, as described in the main results section. In all three steps, the training aims to reduce the mean square error between the input and the reconstructed gene expression vector. Model parameters are updated using Adam optimizer.

*NetActivity* enables adding additional hidden layers in the encoder or decoder layers. A hidden layer added to the encoder would represent multiple scores for the gene sets, and each neuron will only be connected to genes in the corresponding gene set. Then, all neurons in the additional hidden layer representing the same gene set will be connected to the corresponding gene set neuron in the output of the encoder. A hidden layer placed in the decoder will be connected to all neurons of the encoder and all neurons of the output. *NetActivity* also enables tuning the hyperparameters of the different training steps: the number of epochs and learning rate of each training step, the batch size, or the activation function for the hidden and output layers.

### Training of the main model

We trained a model based on GO (Gene Ontology) biological processes^17,18^ and KEGG pathways^19–21^ using the 9,662 samples from GTEx v8^22^. Count and phenotype data were downloaded using *recount2*^47^ R package and count data was transformed using Variance Stabilizing Transformation (VST) from *DESeq2*^48^ R package. Next, we selected the 19,423 coding genes, based on GENCODE v33 annotation^49^, and standardized the data at the gene level (each gene has a mean expression of 0 and a standard deviation of 1). We used 80% of the samples for training and 20% for validation, preserving this proportion for the different GTEx tissues (brain tissues were considered as one tissue). In total, 7,729 samples were used for training and 1,933 samples for validation. Hyperparameters used for training can be found in Sup Table 10.

We first trained a model including all the GO terms and a selection of KEGG terms. We removed KEGG terms representing collections, i.e., groups of genes that do not have a common function (e.g., hsa02010, which contains a collection of ABC transporters), or terms that contain different independent pathways (e.g., hsa04742). We only selected gene sets with more than 10 genes, leading to an initial model with 6,915 gene sets (all GOs + KEGGs model), 6,727 biological processes, and 188 manually filtered KEGG pathways. We trained this model using 6 different weights initializations and computed the Spearman’s rank correlation coefficient between the GSAS for each pair of initializations, using all GTEx samples (train and test samples). We defined replicability as the minimum correlation in absolute value between two initializations. We use the absolute value due to the arbitrariness of the GSAS sign. Thus, a replicability of 1 means that the GSAS had the same values (ignoring the sign) in the 6 initializations, while a replicability of 0 means that in at least two initializations, the correlation between the GSAS was 0.

Next, we removed gene sets with more than 30 genes (Sup Figure 15) and gene sets sharing more than 50% of genes with other gene sets. We retrained the model and evaluated the replicability. Finally, we removed gene sets with replicability < 0.7 and retrained the model. The resulting model, with 1,518 gene sets (Sup Table 1), 1,485 from GO (Sup Table 2), and 33 from KEGG (Sup Table 3), was the main model in the subsequent analyses.

#### Other network configurations

We evaluated the performance of networks with additional hidden layers, maintaining the output of the encoder with 1,518 gene sets. Three additional architectures were tested: (a) Gene Set + Dense - where an additional hidden layer of 1,000 neurons was added at the decoder; (b) Dense + Gene Set - where an additional hidden layer of 10 neurons per gene set was added at the encoder; and (c) Dense + Gene Set + Dense - which combines (a) and (b).

We also tested the effect of adding a dropout layer in the decoder in Step 3 of the training. We trained two models: in “Whole training + dropout” we used the same training as the main model while in “Step 1 + step 3 + dropout”, step 2 of training was not run. In both cases, the dropout rate was set to 0.3, while the remaining hyperparameters were those used for the main model (Sup Table 10).

#### Training steps

We evaluated how the different training steps affected the gene set activity scores. To this end, we computed the replicability obtained when training a model with the initial gene sets (6,915 gene sets) or with the final gene sets (1,518 gene sets) using only step 1 or only steps 1 and 3. We used the same hyperparameters for training these models as for the main model (Sup Table 10).

Next, we assessed whether *NetActivity* gave more relevance to more biologically relevant genes. To compute *NetActivity* gene relevance, we computed the marginal gene relevance for each gene set as the weight of each gene divided by the sum of the weights of all genes in the gene set. Weights were transformed to the absolute value before this computation. Then, *NetActivity* gene relevance was computed as the sum of marginal gene relevance across all the gene sets a gene is included into. The association between *NetActivity* gene relevance and the number of diseases per gene or pLI was run using a linear regression adjusted for the number of gene sets a gene is included into. For the regressions, the number of diseases per gene was log10 transformed adding a 0.1 offset, while pLI values were logit transformed.

Next, we explored how the model defined the gene weights at different training steps. To this end, we extracted the weights of the hsa00430 KEGG pathway in the models fully trained or trained up to step 1. We used the 6 models trained with different weights initialization to assess the model stability, while we used the main model and the first model of step 1 training to perform detailed comparisons of weights and gene set activity scores. Finally, we used GTEx VST transformed data to compute a Principal Component Analysis (PCA) of hsa00430 genes and the Pearson’s correlation between hsa00430 genes.

#### Gene set representativity

We evaluated whether the gene set activity scores contained the information present in the full gene expression matrix. First, we computed the variance of the original gene expression matrix explained by the GSAS in GTEx and TCGA samples using redundancy analysis from the *vegan* R package (https://CRAN.R-project.org/package=vegan). TCGA RNA-seq data was downloaded using *TCGAbiolinks*^50–52^ R package and the effect of the sequencing center was removed with *ComBat-seq*^53^ R package, protecting for project id and sample type (tumor vs normal). Data was transformed using VST from *DESeq2*. The 19,423 genes included in the main model were selected and the data was standardized at the gene level.

Second, we assessed the correlation between the Principal Components (PCs) of the original gene expression matrix and the GSAS in each TCGA project and an IBD cohort (GEO ID: GSE57945)^54,55^. In TCGA, data was standardized at the gene level independently in each project. GSE57945 consisted of an RNA-seq cohort of Intestinal Bowel Disease (IBD), comprising 41 controls, 213 individuals with Crohn’s disease (CD), and 60 with Ulcerative Colitis (UC). Data was downloaded using the *recount2*^47^ R package and transformed with VST. We selected the 19,423 genes in the main model and standardized the data at the gene level. Gene set activity scores were computed with *NetActivity* in each TCGA project and GSE57945. For evaluating the gene set representativity, we run PCA in the original gene matrix and the GSAS matrix of each TCGA project and the IBD cohort. We computed the variability of the first ten GSAS in the original gene expression matrix using Redundancy Analysis from *vegan* R package. We further explored the gene set representativity in TCGA prostate cancer samples (PRAD) and the IBD cohort, by computing the Pearson’s correlation between the top ten PCs of the original gene matrix and the top ten PCs of the gene activity scores.

### Comparison of NetActivity with other gene set projection methods

We compared the performance of *NetActivity* with two other gene set projection methods: GSVA and hipathia. GSVA scores were computed using the *GSVA* R package^9^ with the same gene sets as in the main model; while Hipathia scores were computed using *hipathia* R package^13^ and the default pathways.

#### Stability of gene set scores definition

To evaluate the stability of gene set scores definition, i.e., whether the gene set scores of a sample are stable to changes in the population, we computed GSAS with *NetActivity*, *GSVA,* and *hipathia*, using all PRAD samples or including only control samples. Then, we defined two measures: (1) score replicability; and (2) definition consistency. Score replicability was defined as the Pearson correlation between the GSAS of control samples when computed using all PRAD samples or only control samples. To compute the definition consistency, we computed, for each gene set, the Pearson correlation between the expression of the genes in the gene set and the GSAS. These correlations were considered as a proxy of the gene set score definition. Thus, we computed the Pearson correlation between the genes’ correlation when computing the GSAS using all PRAD samples or only control samples.

Next, we evaluated whether the gene set score definitions were consistent in different datasets. We used three different datasets: TCGA-PRAD, TCGA-BRCA, and GEO-PRAD. TCGA-PRAD consisted of the 334 prostate cancer samples from TCGA that contained Gleason information^56^. We classified the 334 samples from PRAD in 88 samples with high Gleason (Gleason > = 8), and 246 with low Gleason (Gleason < 8). TCGA-BRCA are 1,066 well-characterized breast cancer samples from TCGA^57^. TCGA-PRAD and TCGA-BRCA were generated with RNA-seq. GEO-PRAD are 1,131 prostate cancer samples from GSE169038^58^ generated with Affymetrix Human Exon 1.0 ST Array. We downloaded GSE169038 with *GEOquery* R package^59^, which consisted of 1,152 prostate cancer samples from European and American ancestries. After discarding samples with a very initial prostate cancer stage (Gleason score < 3), we had 1,131 samples, 115 with high Gleason (>= 8), and 1,016 with low Gleason (< 8). To prepare the data for *NetActivity*, we mapped GSE169038 features to ENSEMBL ids, using the platform annotation. If multiple features were mapped to the same ENSEMBL id, the feature with more probes was selected. *NetActivity* was then used to standardize the data and compute the scores. We computed GSAS with *NetActivity*, *GSVA,* and *hipathia* in TCGA-PRAD, TCGA-BRCA, and GEO-PRAD. We compared the GSAS definition between TCGA-PRAD and GSE169038 (same tissue, different technology - RNA-seq vs Affymetrix Human Exon 1.0 ST Array), and between TCGA-PRAD and TCGA-BRCA (different tissue, same technology).

#### Replicability

We evaluated whether we obtained the same results when performing the same analysis for different projection methods (*NetActivity, GSVA,* and *hipathia*) and individual gene expression in different datasets of the same disease. We compared the gene expression between prostate samples with low or high Gleason scores, a histological marker of tumor progression. We computed the gene set scores using *NetActivity*, GSVA, and hipathia, based on VST values for TCGA-PRAD and the original gene expression values for GEO-PRAD.

All models in PRAD were adjusted for tumor subtype, age, and ancestry, while models in GSE169038 were adjusted for ancestry and decipher classification. Individual gene differential expression analysis was run using raw counts and *DEseq2* R package for PRAD and original gene expression values and *limma*^60^ R package for GEO-PRAD. In both cases, we used all available genes for differential expression analysis and false discovery rate (FDR) computation. Then, GEO-PRAD features were mapped to ENSEMBL ids, and genes annotated to the same ENSEMBL id were selected. The differential expression analysis for *NetActivity* and GSVA gene set scores was run using *limma* R package. For *hipathia* gene set scores, the differential expression analysis was run using the Wilcoxon test implemented in *hipathia* R package. Finally, we computed the correlation between the differential expression estimates in both datasets: (1) genes - log2FC from *DESeq2* and logFC in limma; (2) *NetActivity* and *GSVA* - logFC in *limma*; and (3) *hipathia* - U-statistic.

### Application of NetActivity

#### Meta-analysis of prostate cancer

We run a meta-analysis of differences in gene expression due to cancer progression in prostate cancer to show the applicability of *NetActivity*. We included nine datasets generated with three platforms: RNA-seq, Illumina HumanHT-12 array, and Affymetrix Human Exon 1.0 ST Array (Sup Table 4).

We preprocessed GSE46691 and GSE21034 CEL files with the rma algorithm from *oligo*^61^ Bioconductor package, summarizing expression values into core genes. Gene probes were mapped to ENSEMBL ids using *huex10sttranscriptcluster.db* Bioconductor package. We normalized count data from GSE183019 and GSE201284 using VST from *DESeq2* R package, while we used normalized gene expression values from GEO for GSE141551, GSE70768, GSE70769. Gene SYMBOL identifiers were converted to ENSEMBL ids with *org.Hs.eg.db* Bioconductor package (https://bioconductor.org/packages/org.Hs.eg.db) in GSE141551, GSE70768, GSE70769, GSE183019 and GSE201284. Pre-processing of TCGA-PRAD and GSE169038 was defined in the previous section. In all datasets, *NetActivity* was used to standardize gene values and to compute GSAS.

In each dataset, we run a differential expression analysis between samples with low (<8) or high (>=8) Gleason scores with *limma*, adjusting for the most relevant covariates in each dataset (Sup Table 4). We regressed out the first 2 surrogate variables from GSE183019, computed with *sva*^62^ Bioconductor package, to correct a clustering of the samples not related to any biological variable. Meta-analysis of effect estimates was run with METAL^63^. To interpret the results, we run a differential expression analysis using the standardized gene values computed in the previous sections, *limma,* and the relevant covariates defined in the previous section.

#### Application to metastatic prostate cancer

We applied *NetActivity* to the RNA-seq data from the PROMOTE study^29,30^ available at dbGAP database with accession number phs001141.v2.p1. PROMOTE study aims to identify the molecular mechanisms associated with abiraterone therapy resistance in metastatic castration-resistant prostate cancer. Samples were taken after 12 weeks of abiraterone acetate/prednisone treatment and the primary outcome was determined by the time-to-treatment-change (TTTC), i.e., the time the patients were under the therapy before they changed to another alternative due to disease relapse or progression. We evaluated the association between the TTTC with the gene expression from RNA-seq data and the activity scores computed from it by *NetActivity*.

We select the 47 samples from bone metastatic tissue, the tissue with the largest number of samples. Original gene Symbols were updated to the newest version based on the HGNG HUGO Gene Nomenclature Committee (https://ftp.ebi.ac.uk/pub/databases/genenames/hgnc/archive/monthly/tsv/hgnc_complete_set_2022-10-01.txt, accessed the 07/10/2022) and mapped to ENSEMBL ids using *AnnotationsDbi* (https://bioconductor.org/packages/AnnotationDbi) and *org.Hs.eg.db* R packages. Count expression data was normalized using VST from *DESeq2* R package, before computing GSAS with *NetActivity*. We explored the effect of log10(TTTC adjusted) (for simplicity from here on referred to as TTTC) on gene expression, using the transformed gene expression matrix or GSAS. Differential gene expression analysis was performed with *DESeq2* R package, while the association of GSAS with TTTC was estimated using a linear model with the *limma* R package. Additionally, we used a linear regression model to estimate the effect of the gene expression of the top gene and the whole set of genes for each of the significantly activated pathways on the TTTC. We calculated the adjusted R2 of these models and compared them with the model computed with the activity scores for those same pathways.

For calculating the differentially expressed genes we used the standard differential expression analysis of *DESeq2* R package using Benjamini and Hochberg’s method for multiple test correction. Overrepresentation enrichment analysis was performed using the function *enrichGO* from the R package *clusterProfiler*^64,65^ R package with size of gene sets to test between 10-5000, using either all genes or just genes included in *NetActivity*.

## Availability of data

Accession codes for public data is referenced in the methods section or in Supplementary Table 4. The code to reproduce the figures, tables and results is deposited in Github (https://github.com/yocra3/NetActivity_paper). A docker container to reproduce the environment used to process the data is available in DockerHub (https://hub.docker.com/r/yocra3/netactivity_analysis). NetActivity framework is available in GitHub (https://github.com/yocra3/NetActivityTrain), while NetActivity model is available in a R/Bioconductor package (https://bioconductor.org/packages/NetActivity).

