## Supplementary Figures for "NetActivity enhances transcriptional signals by combining gene expression into robust gene set activity scores through interpretable autoencoders"

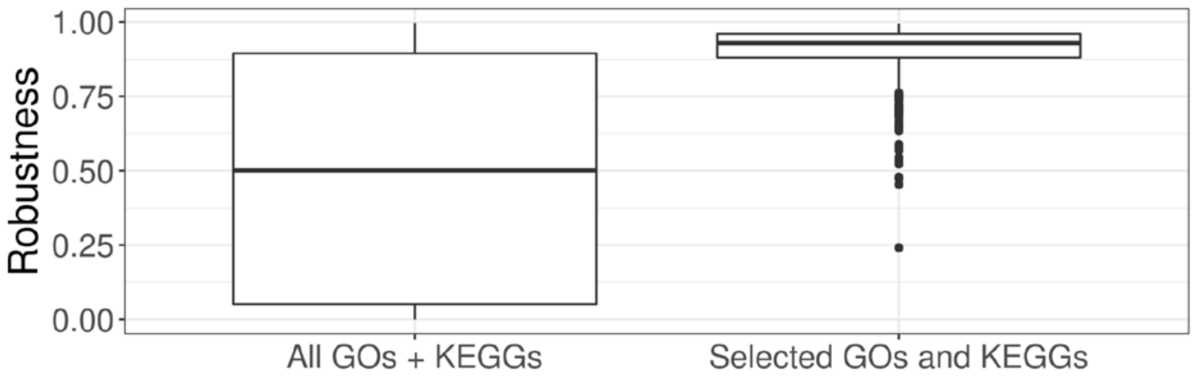

**Sup Fig 1: Robustness due to gene set selection.** Higher robustness means higher stability of the GSAS across different initialization weights. All GOs + KEGGs: model trained using 6,915 GO terms and KEGG pathways. Selected GOs and KEGGs: model trained using the selected 1,518 GO terms and KEGG pathways. Both models were trained using the three-step approach and included a gene set encoder and decoder with a single layer.

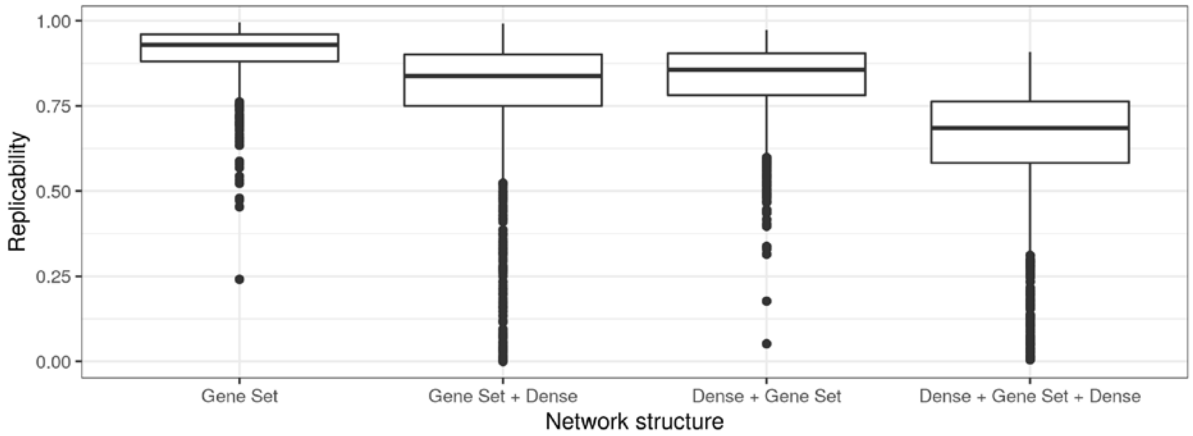

**Sup Fig 2: Replicability in different network structures**. Higher replicability means higher stability of the GSAS across different initialization weights. All models were run with the selected 1,518 pathways. Gene Set: main model, with only the gene set activity layer. Gene Set + Dense: this model includes an additional layer in the decoder between the gene set activity layer and the output layer. Dense + Gene Set: this model includes a shallow connected layer in the encoder between the input layer and the gene set activity layer. Dense + Gene Set + Dense: this model includes the two additional layers of the previous two models.

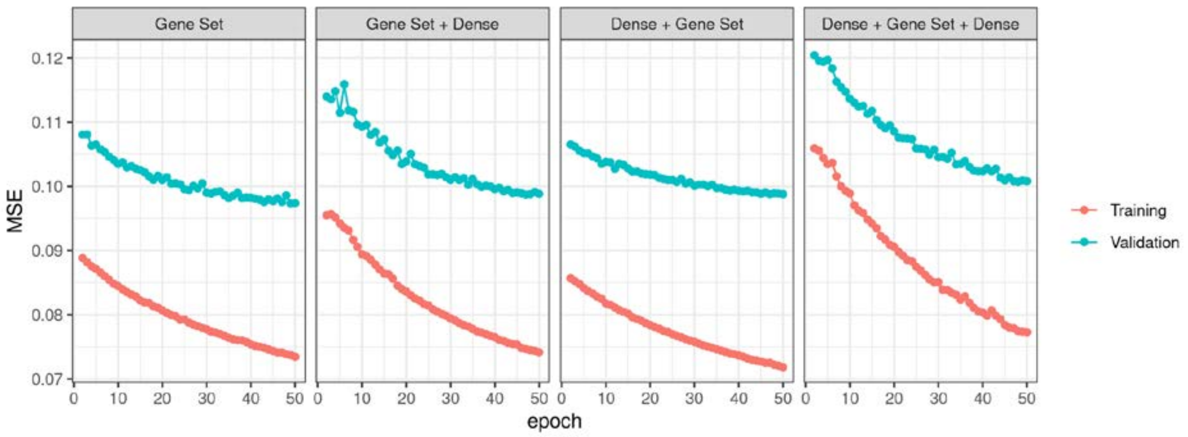

**Sup Fig 3: Training performance in different network configurations**. A lower Mean Squared Error (MSE) means a more accurate reconstruction of the original gene expression values from the GSAS. MSE was computed for training samples (red) and for validation samples (blue). All models were run with the selected 1,518 gene sets. Gene Set: main model, with only the gene set activity layer. Gene Set + Dense: this model includes an additional layer in the decoder. Dense + Gene Set: this model includes an additional shallow connected layer in the encoder. Dense + Gene Set + Dense: this model includes an additional layer in the encoder and decoders. MSE for the first epoch is not shown to reduce the y-axis range.

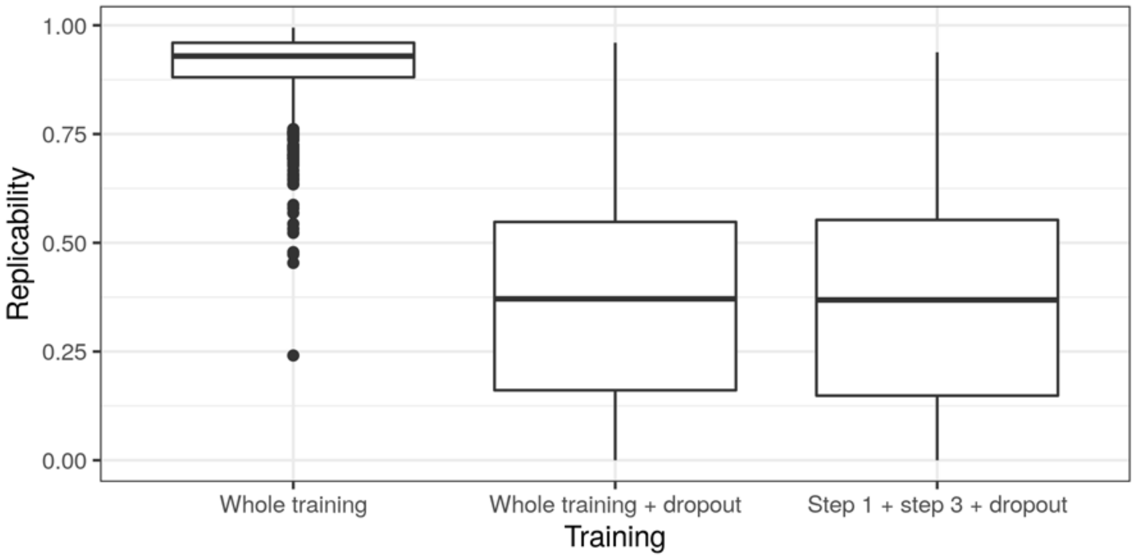

**Sup Fig 4: Replicability in different training strategies**. Higher replicability means a higher stability of the GSAS across different initialization weights. All models were run with the selected 1,518 gene sets. Whole training: main model without dropout. Whole training + dropout: whole training with dropout added in step 3 before the output layer. Step 1 + step 3 + dropout: same as Whole training + dropout, but without step 2 (freezing the weights of the encoder).

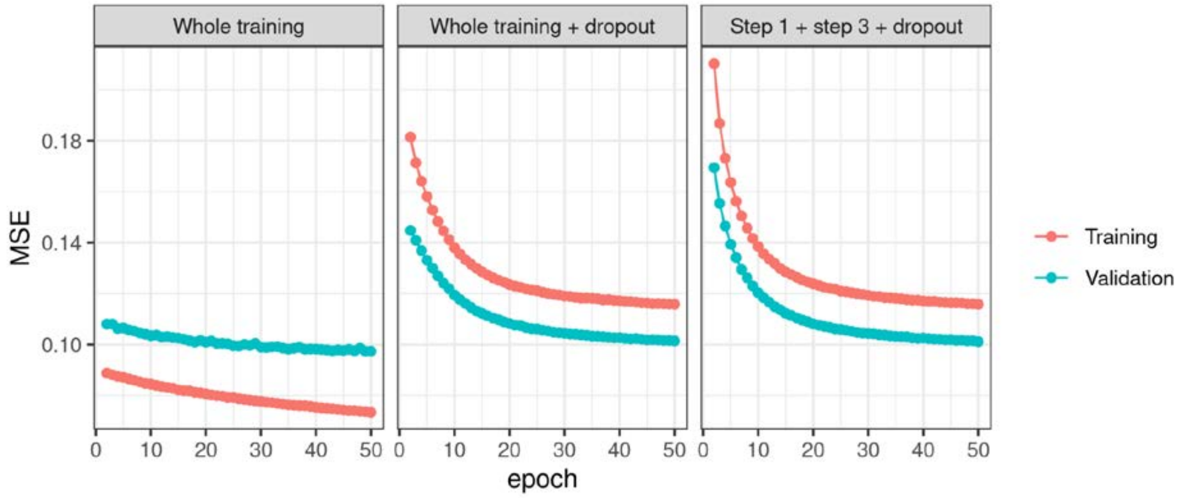

**Sup Fig 5: Training performance in different training configurations.** A lower Mean Square Error (MSE) means a more accurate reconstruction of the original gene values from the GSAS. MSE was computed for training samples (red) and for validation samples (blue). All models were run with the selected 1,518 gene sets. Whole training: main model without dropout. Whole training + dropout: whole training with dropout added in step 3 before the output layer. Step 1 + step 3 + dropout: same as Whole training + dropout, but without step 2 (freezing the weights of the gene set activity layer).

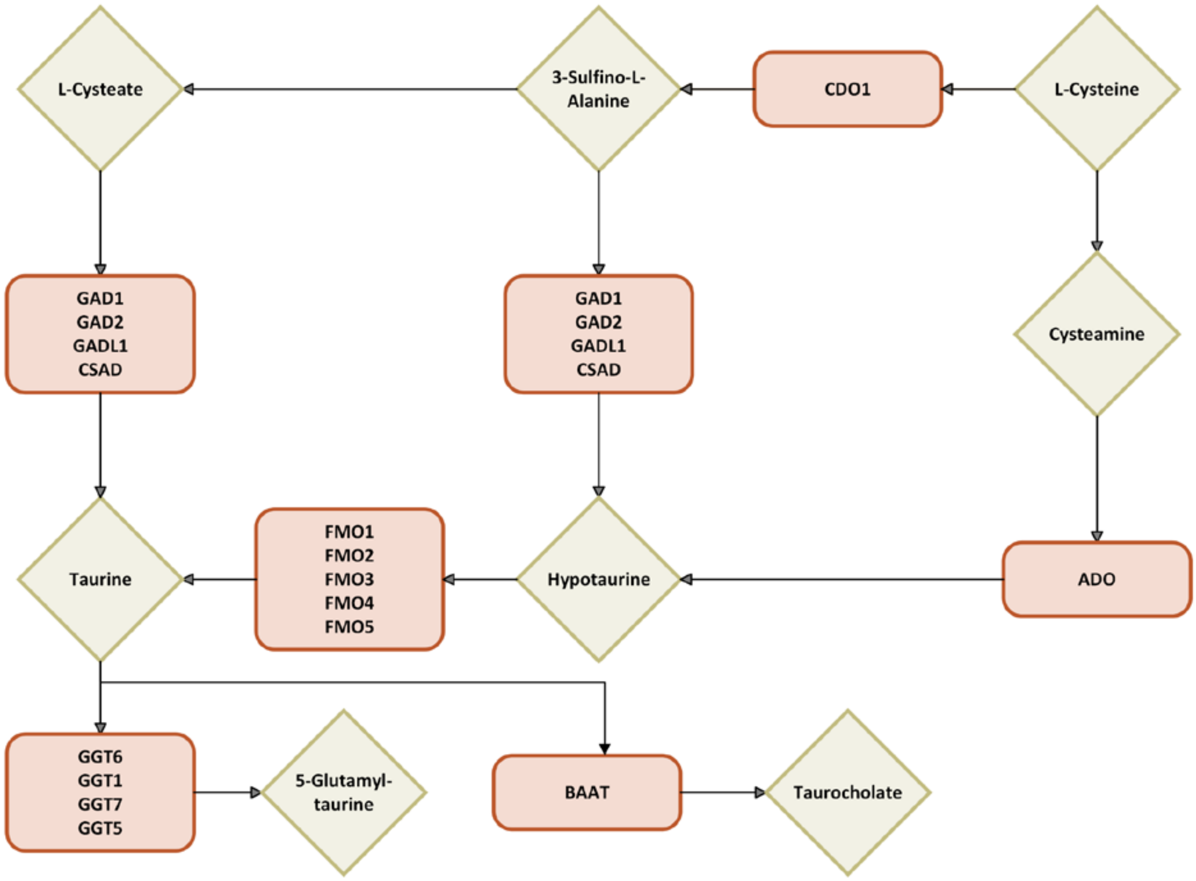

**Sup Fig 6: hsa00430 KEGG pathway**. This pathway represents the taurine and hypotaurine metabolism with human genes. Green diamonds represent compounds, while orange rectangle genes. Multiple genes in the same rectangle represent multiple genes that can catalyze the same reaction.

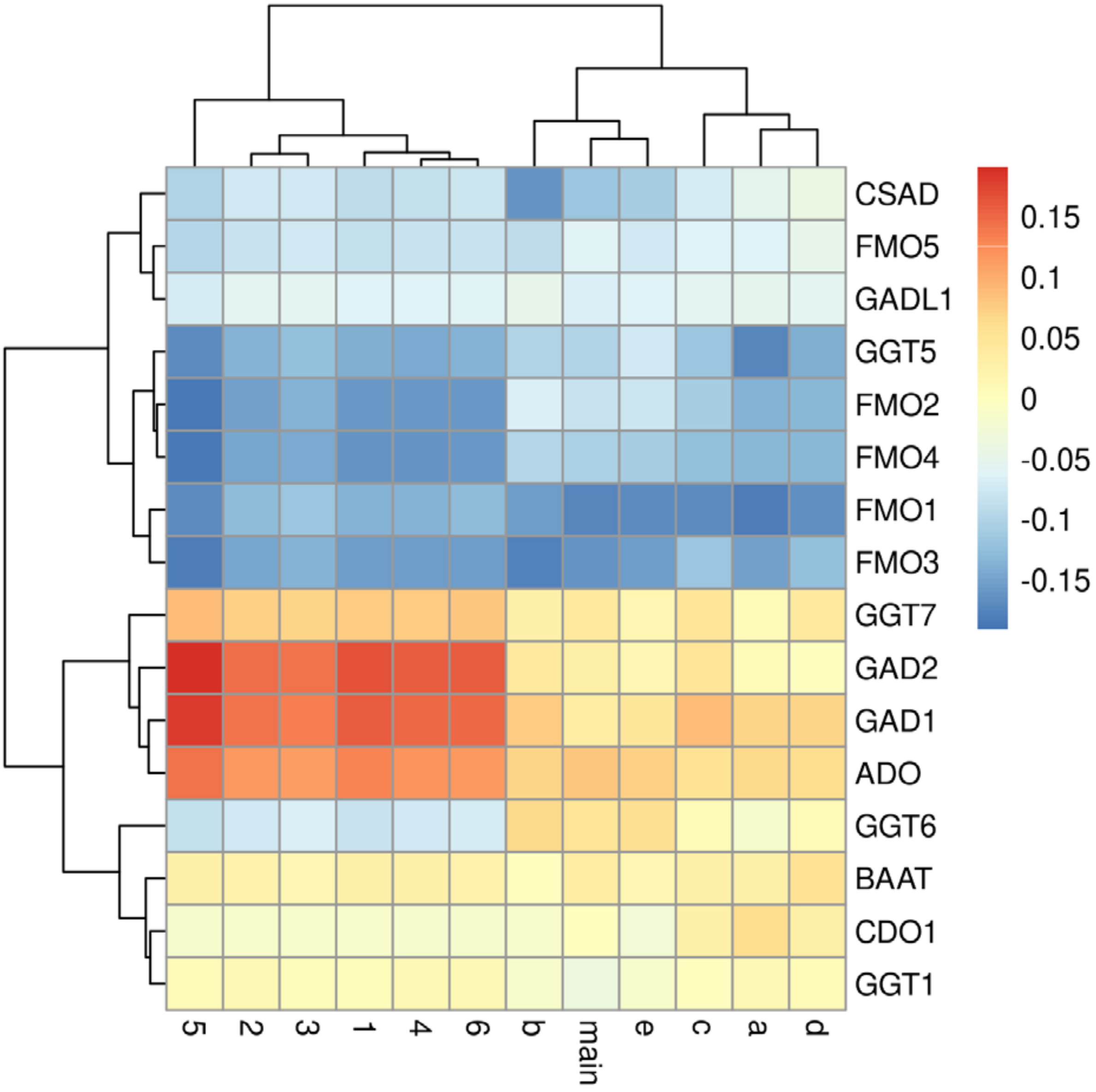

**Sup Fig 7: Weights defined by NetActivity for hsa00430 KEGG pathway**. Columns represent different model initializations: in numbers (1-6), models trained only until step 1; in letters (a-e) models with full training; main represents the main model. The sign of the weights of models a, c and d has been changed to ease the comparison with the other models. Higher magnitude weights are represented with darker colours.

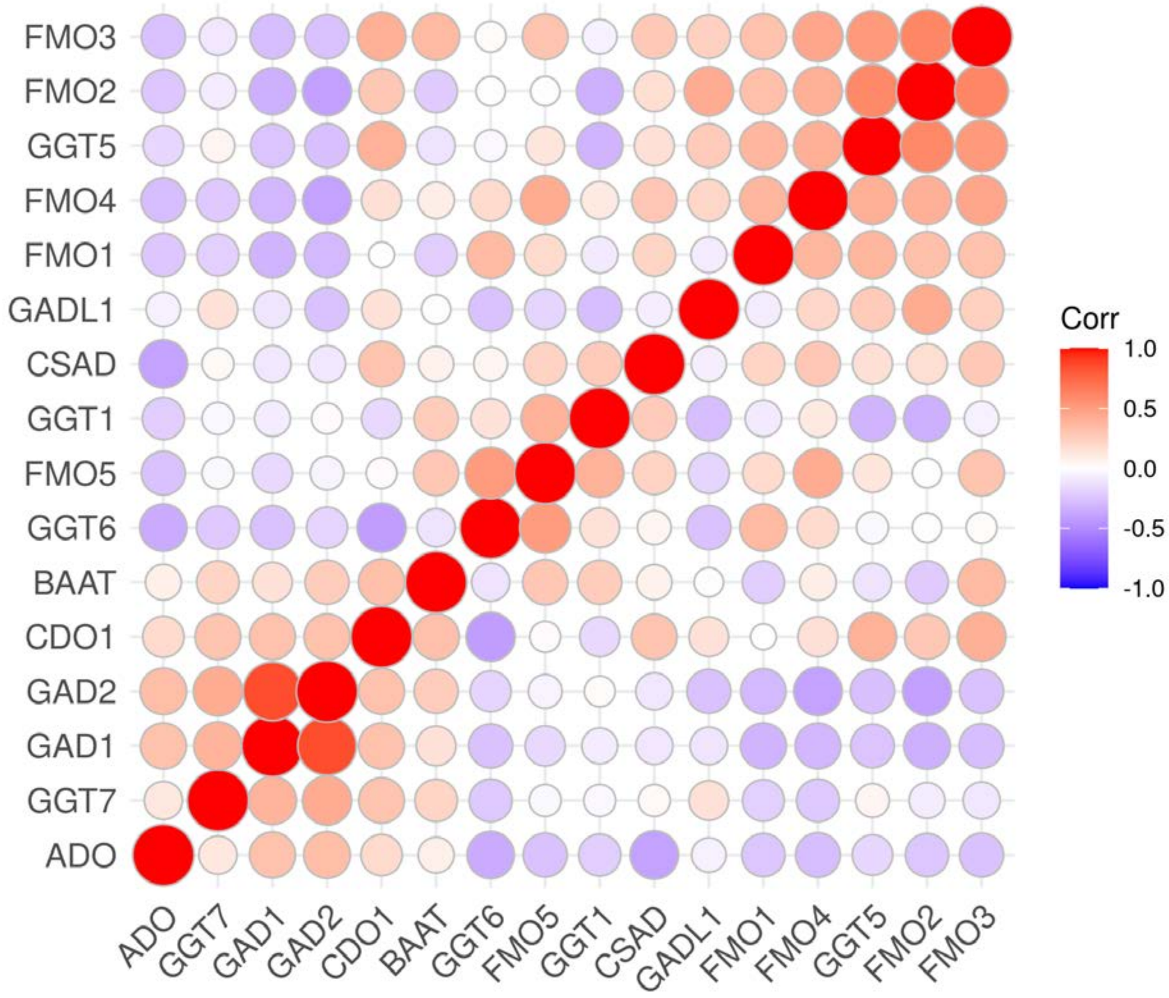

**Sup Fig 8: Pearson correlation of genes in hsa00430 KEGG pathway in GTEx**. Positive correlations are shown in red and negative in blue. Genes are grouped based on a hierarchical clustering to highlight clusters of correlated genes.

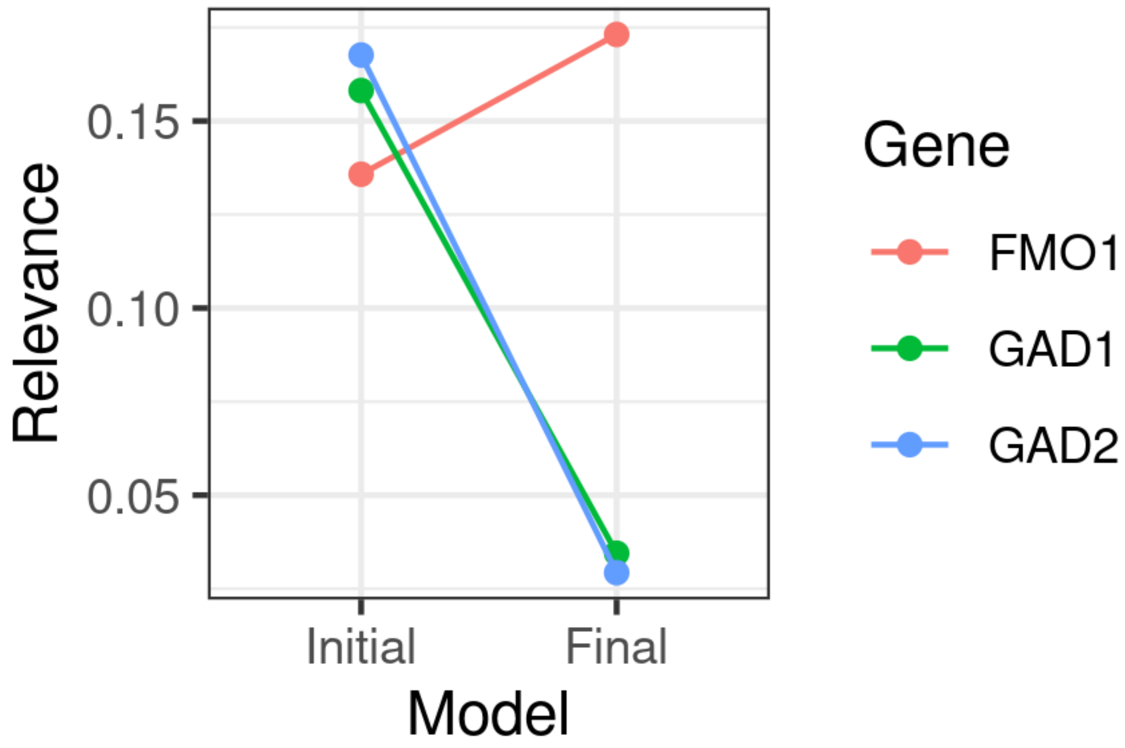

**Sup Fig 9: Change of weights in hsa00430 KEGG pathway**. Initial is the first model trained until step 1. Final is the main model. Only the three most relevant genes are shown. Relevance are the weights represented in absolute value.

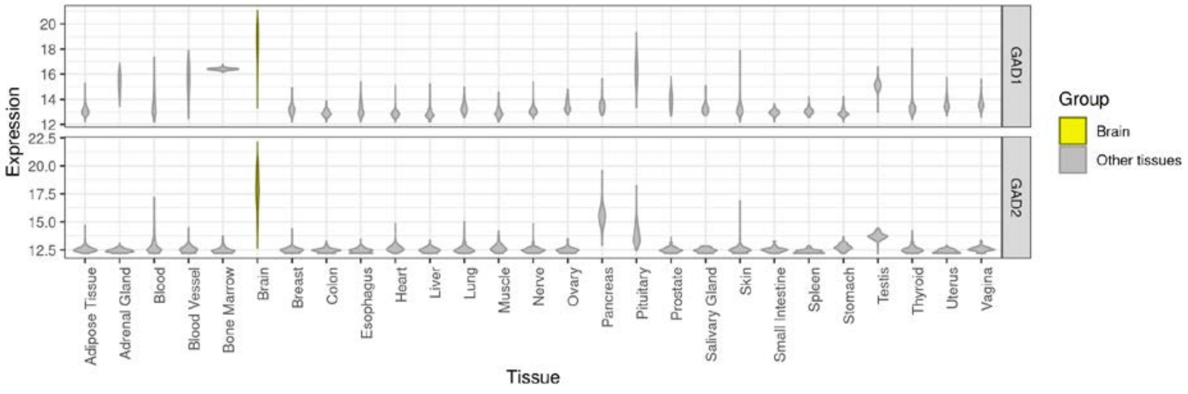

**Sup Fig 10: Gene expression values for GAD1 and GAD2 in GTEx per tissue.** Expression values correspond to *DESeq2* VST (Variance Stabilization Transformation) values. In yellow, gene expression values in brain tissues.

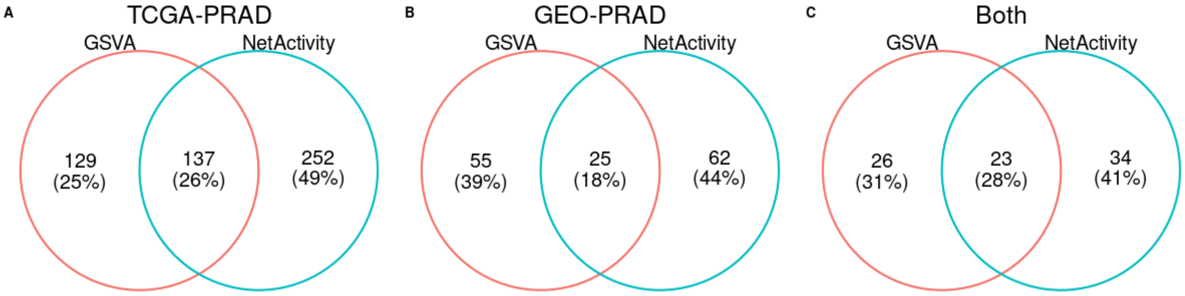

**Sup Fig 11: Overlap between the differentially expressed gene sets using *GSVA* and *NetActivity***. A: differentially expressed gene sets in PRAD samples from TCGA (TCGA-PRAD). B: differentially expressed gene sets in GSE169038 (GEO-PRAD). C: gene sets identified as differentially expressed in TCGA-PRAD and GEO-PRAD.

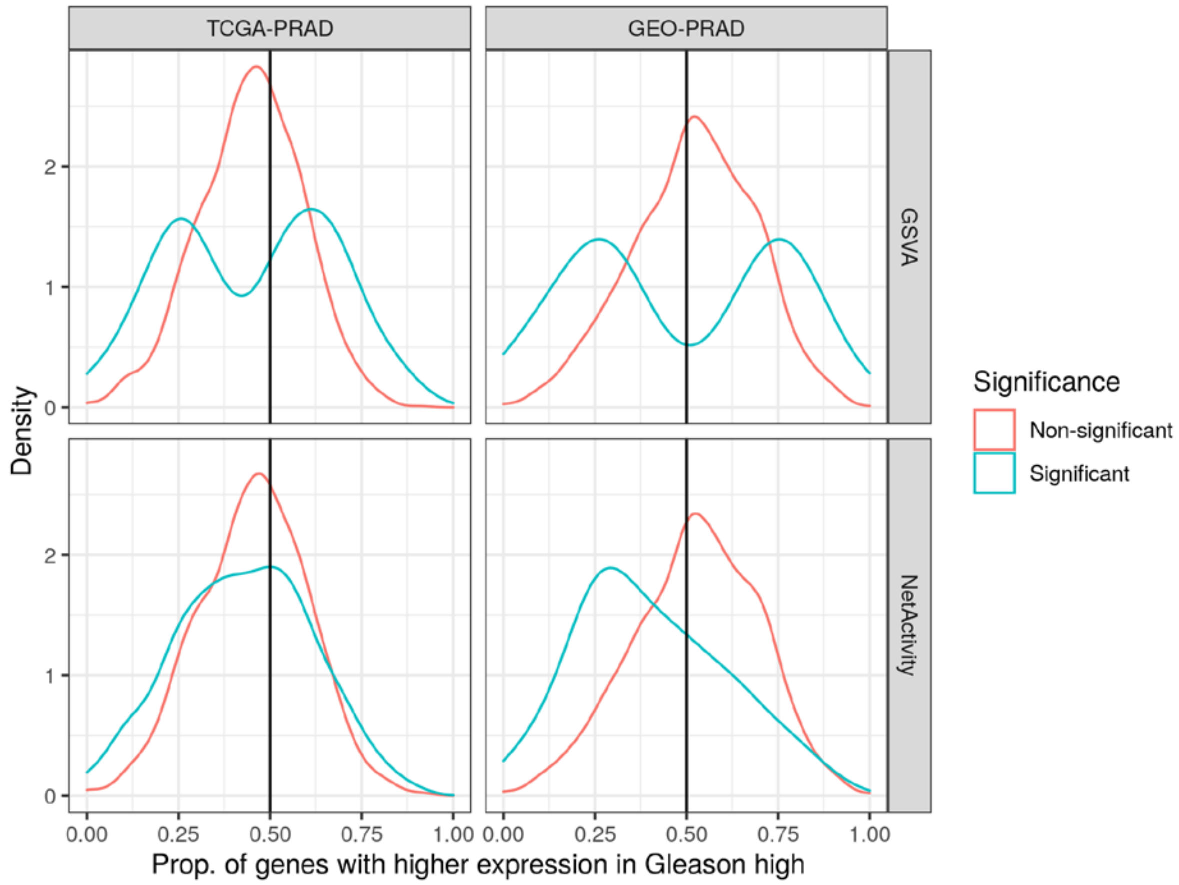

**Sup Fig 12: Distribution of the proportion of gene set genes with higher expression in high Gleason individuals**. For each gene set, we computed the proportion of genes that had a higher expression in high Gleason individuals, without considering the statistical significance. Gene sets were grouped on whether they were significant (FDR < 0.05) or not (FDR > 0.05), on a given combination of dataset (TCGA-PRAD or GEO-PRAD) and gene set projection method (*NetActivity* or *GSVA*). TCGA-PRAD: PRAD samples with Gleason information. GEO-PRAD: samples from GSE169038.

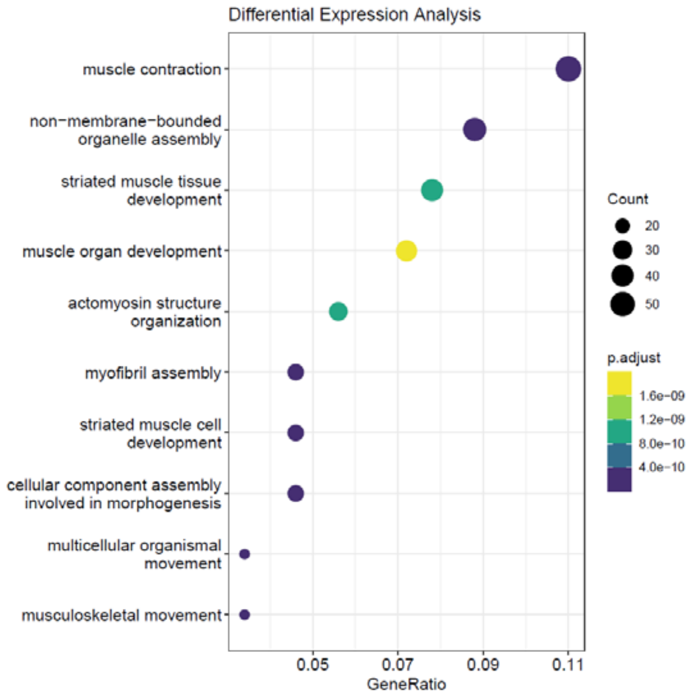

**Sup Fig 13: Top 10 enriched GO biological process terms in samples responding to abiraterone in the PROMOTE study**. Enrichment was computed based on the 592 significantly differentially expressed (DE) genes of PROMOTE samples (FDR < 0.05). X-axis represents the ratio of DE genes present in the GO term over the total number of genes in the gene set and dots' sizes represent the total number of DE genes coincident with the GO term genes. p-adjust is calculated with Benjamini & Hochberg correction that accounts for the false discovery rate (FDR).

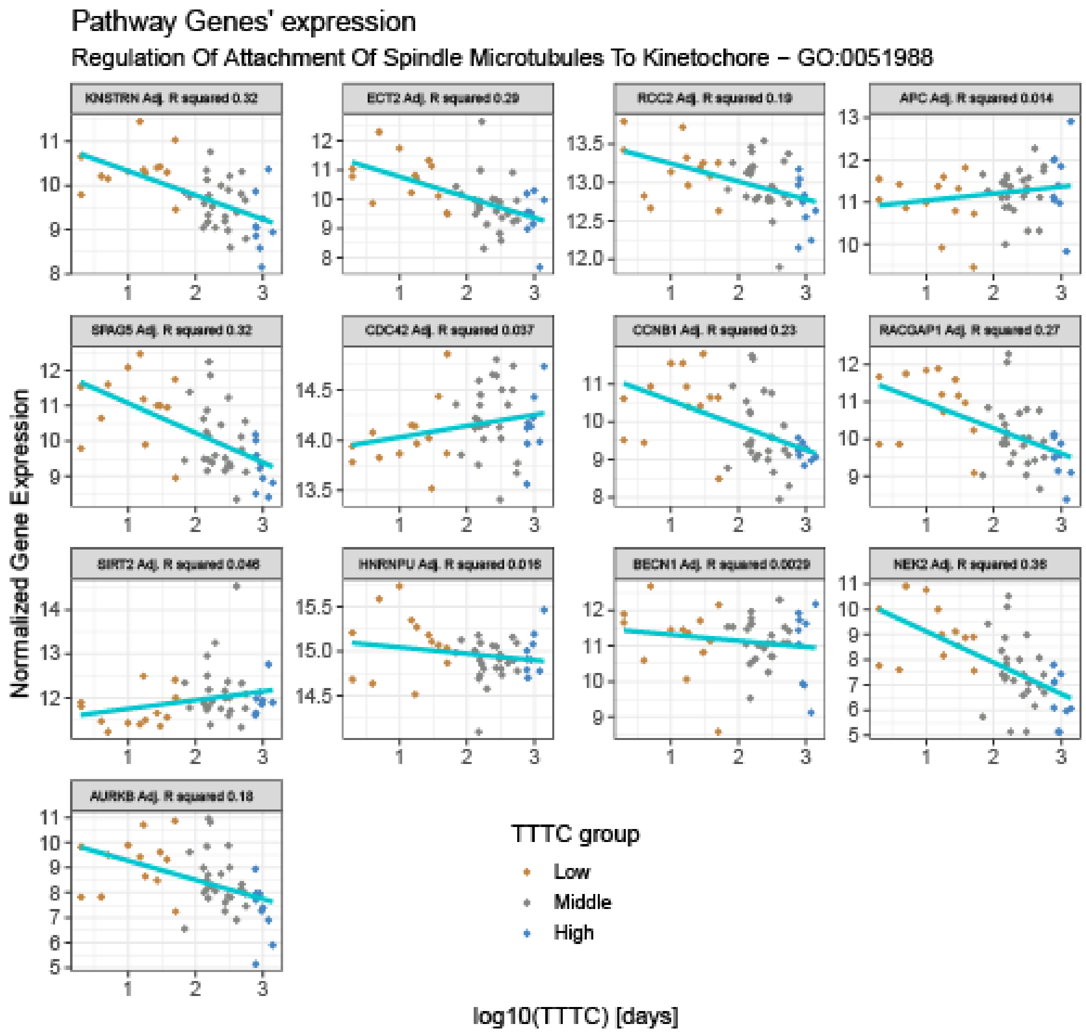

**Sup Fig 14: Expression of genes of the top gene set in PROMOTE data**. GO:0051988, regulation of attachment of spindle microtubules to kinetochore, was the gene set with the highest association with abiraterone treatment. Expression of genes in GO:0051988 was normalized using Variance Stabilizing Transformation (VST) from *DESeq2*. Samples were colored based on their TTTC (Time To Treatment Change): in orange, samples in the lowest TTTC quantile (low); in grey, samples in the second and third TTTC quantiles (Middle); in blue, samples in the top TTTC quantile (High). The regression line shows the association between the log10 of TTTC and the expression of each gene. Each box contains the proportion of variance of the TTTC explained by the gene expression (Adj.R squared).

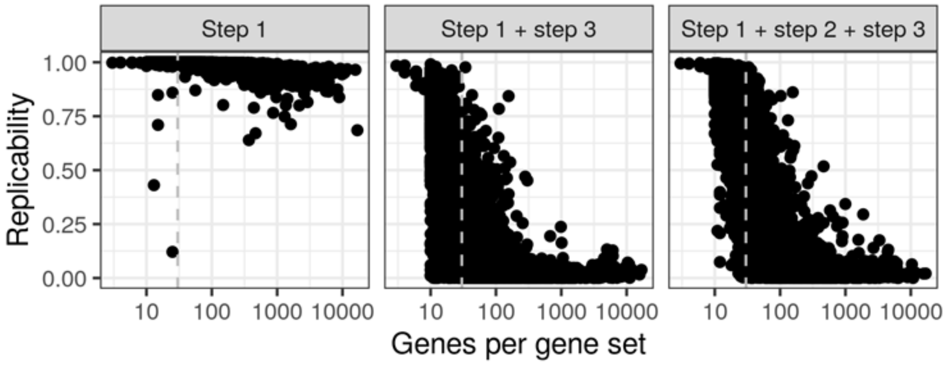

**Sup Fig 15: Effect of gene set size on GSAS replicability.** Higher replicability means higher stability of the GSAS across different initialization weights. The vertical dashed line is set at 30 genes per gene set, the threshold used to filter gene sets. Step 1: only the training of a dense autoencoder per gene set was performed. Step 1 + step 3: initialize the encoder with the weights from step 1 and then train the whole shallow sparsely connected autoencoder (encoder and decoder). Step 1 + step 2 + step 3: freeze weights of gene set activation layer (step 1) and train the decoder for some epochs (step 2), before training the shallow sparsely connected autoencoder (step 3).
